# Analysis tools to quantify dissemination of pathology in zebrafish larvae

**DOI:** 10.1101/639633

**Authors:** David R. Stirling, Oniz Suleyman, Eliza Gil, Philip M. Elks, Vincenzo Torraca, Mahdad Noursadeghi, Gillian S. Tomlinson

## Abstract

We used the zebrafish larval *Mycobacterium marinum* infection model of tuberculosis to develop image analysis software called QuantiFish, to enable rapid, objective quantitation of bacterial load and dissemination. Our analysis tools provide novel outcome measures of infection; the proportional distribution of bacteria between sites and their spatial distribution throughout the host. We anticipate the broader application of these parameters to other zebrafish models of infection and metastatic cancer.

---

Zebrafish are increasingly being used to address biological questions in the life sciences. Advantages of this model include genetic tractability, faithful representation of mammalian systems and optical transparency during the larval stages^1^. The availability of transgenic lines with fluorescent immune cell lineages and fluorescent protein expressing microbes provide unparalleled opportunities for quantitative in vivo imaging, particularly in relation to host-pathogen interactions^2^. Currently, the outcome of zebrafish larval infection models is almost exclusively measured by quantitative imaging of fluorescent pathogens as a surrogate for total pathogen burden. This approach fails to consider pathogen dissemination as an alternative measure of control of infection. In order to address this limitation in current practice, we have developed new, open source image analysis software called QuantiFish. This enables rapid and objective quantitation of pathogen load and of dissemination, using a range of parameters. We tested this software using *Mycobacterium marinum* (Mm) infection of zebrafish larvae, widely used to model human tuberculosis (TB)^3^. We anticipate that the software will also be applicable to other zebrafish models of infection and metastatic cancer.

To develop our dissemination analysis tools we used images of Mm infected larvae (Supplementary figure 1a) concordantly classified by two independent investigators as examples of minimally, moderately or widely disseminated infection (Supplementary figure 1b). Images were analysed with QuantiFish, which provides a simple interface for rapid, sensitive quantitation of total fluorescence and signal from individual fluorescent foci (Supplementary figure 2). Total bacterial load (integrated fluorescence intensity) and numbers of separate foci of bacteria detected using QuantiFish were significantly higher in fish with more widely disseminated infection (Supplementary figure 3a, d). Comparable results were obtained using a published ImageJ macro^4^ (Supplementary figure 3b, e) and found to be very highly correlated with those generated using QuantiFish (Supplementary figure 3c, f).

However, our software represents a significant advance, both because of its intuitive interface and more importantly, its ability to generate detailed information about the characteristics of each individual fluorescent focus. We exploited this to develop new analysis tools to evaluate features of bacterial dissemination which are not possible to assess using existing strategies that measure area or intensity of bacterial fluorescence or count the total number of fluorescent foci without any regard to their spatial distribution^4^. We conceptualised the limitations of current methods in schematic representations of different distributions of fluorescent foci (Figure 1a). The first example (Figure 1ai) compares differences in the distribution of infection that are measured by current methods to quantify the total fluorescence or the number of fluorescent foci. However, neither measure distinguishes differences in the proportion of fluorescence in different foci, for example if 90% of the bacteria are contained within one of two foci, or equally distributed between two foci (Figure 1aii). Nor can they discriminate localised dissemination of infection from distant dissemination of infection (Figure 1aiii). To address these limitations, we propose four measures of bacterial dissemination. First, the number of fluorescent bacterial foci that are responsible for 50% of the total fluorescence; second the number of predefined grid zones that contain the centre point of a bacterial focus (Figure 1bi); third the area of a polygon containing the centre points of all foci (Figure 1bii) and fourth the maximum distance between any two foci (Figure 1biii).

**Fig. 1.**
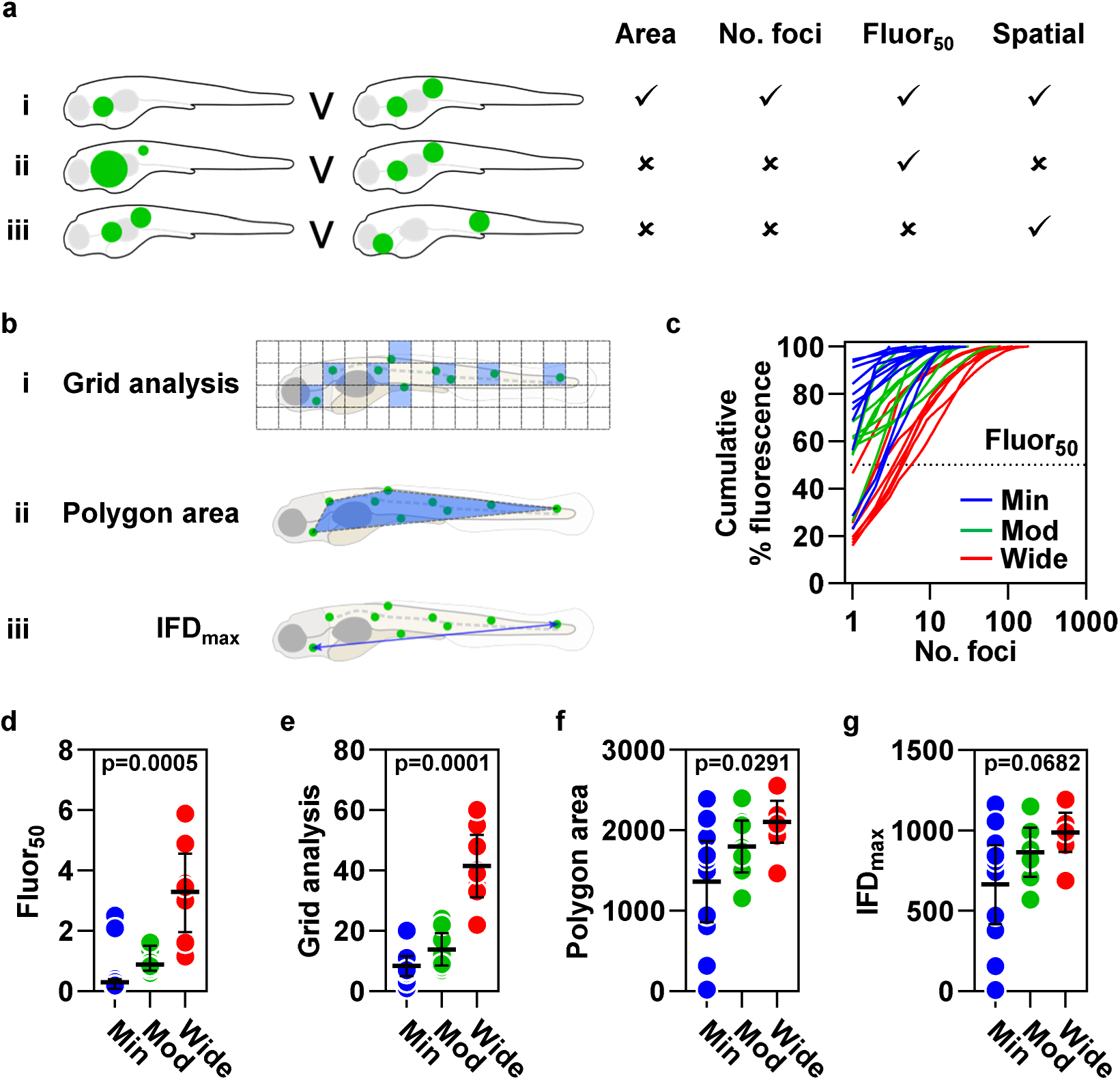
Analysis tools to quantify dissemination of mycobacterial infection. **a**, Limitations of existing outcome measurements of infection, illustrated in schematic diagrams representing different distributions of fluorescent bacterial foci (green). Area of fluorescent signal and number of foci distinguish some distributions of infection, **ai**, but not others **aii-iii**. We propose fluor_50_, the number of foci that contribute 50% of the total fluorescence, to distinguish differences in the proportional distribution of the total burden of pathology at each site, **aii**, and parameters that quantify the spatial distribution of foci to differentiate localised dissemination from distant dissemination, **aiii. bi**, Grid analysis divides the image into an array of squares, then quantifies the number of grid zones containing the centre point of ≥1 foci (highlighted blue). **bii**, The area of a polygon (highlighted blue) encompassing the centre points of all foci. **biii**, The maximum distance between the centre points of any two foci (IFD_max_). **c-g**, Quantitation of bacterial dissemination in zebrafish larvae 4 days post infection (dpi) with 200-400 cfu *Mycobacterium marinum* expressing mWasabi, classified as having minimally, moderately or widely disseminated infection (n=11, 8 and 8, respectively). **c**, The relationship between cumulative percentage fluorescence and the number of foci which generate this signal. **d**, Fluor_50_, calculated by interpolation of the data shown in **c**. Spatial measurements of dissemination, **e**, grid analysis, **f**, polygon area and **g**, IFD_max_. Lines **c**, and data points, **d-g**, represent individual zebrafish larvae. Lines and error bars, **d-g**, represent median ± IQR. p values were derived from Kruskal-Wallis tests with Benjamini, Krieger and Yekutieli correction for multiple comparisons.

We reasoned that in less disseminated infection a higher proportion of the bacterial fluorescence would be localised in fewer foci. This was demonstrated by plotting the percentage cumulative fluorescence in each fish as a function of the number of fluorescent foci (Figure 1c). A summary statistic for each fish was then derived by interpolation of these data to estimate the number of fluorescent foci responsible for 50% of the total fluorescence (fluor_50_). This measure reliably distinguished different groups of fish independently classified as having minimally, moderately or widely disseminated infection (Figure 1d).

We also sought to develop strategies to assess the spatial distribution of fluorescent foci, on the premise that bacterial foci distributed over a broader area represent more highly disseminated infection, indicative of more severe disease. Our first approach, termed “grid analysis”, quantifies the number of user-defined grid zones that contain the centre point of one or more foci of fluorescence. Multiple fluorescent foci located within a single zone are counted as one positive grid section, emphasizing larger differences in spatial distribution than simply counting the number of foci (Figure 1bi). Our second approach uses the co-ordinates that describe the locations of the centre points of individual fluorescent foci to compute the area of the smallest convex polygon that contains all these points (Figure 1bii). Our third strategy extends this approach by using the points that define the boundaries of this polygon to calculate the maximum distance between any two fluorescent foci (IFD_max_) (Figure 1biii). These three measures were each concordant with the investigators classification of the degree of dissemination. The spatial distribution of bacteria was highest in widely disseminated infection, intermediate in moderately disseminated infection and lowest in minimally disseminated infection. Both grid analysis and polygon area showed significant differences between minimally, moderated and widely disseminated infection (Figure 1e-g).

We then tested the hypothesis that infection with higher doses of Mm would lead to greater dissemination. First, we demonstrated the expected incremental increase in bacterial burden in zebrafish larvae infected with a dose titration of Mm (25, 100 and 400 colony forming units (cfu)), using both QuantiFish and ImageJ, which provided highly correlated results (Supplementary figure 4a-c). Treatment with the anti-mycobacterial drug, isoniazid, significantly diminished detected fluorescence, consistent with control of infection (Supplementary figure 4a, b). Escalation of the inoculum dose also led to higher numbers of fluorescent foci, and treatment with isoniazid was associated with significantly fewer foci (Supplementary figure 4d-f). Next, we applied our dissemination analysis tools to quantify the proportional distribution of the total bacterial load between sites and the spatial distribution of bacteria within each fish. Higher inoculum dose was associated with significantly increased dissemination, demonstrated by smaller proportions of the total fluorescence divided between larger numbers of foci (Figure 2a, b) and by the three measurements that reflect the spatial distribution of bacteria (Figure 2c-e). We interpret these data as evidence that inoculum dose directly influences both the spatial dispersion of bacteria, and the ability of the host to predominantly contain bacterial growth to fewer sites, reflected by the fluor_50_ parameter.

**Fig. 2.**
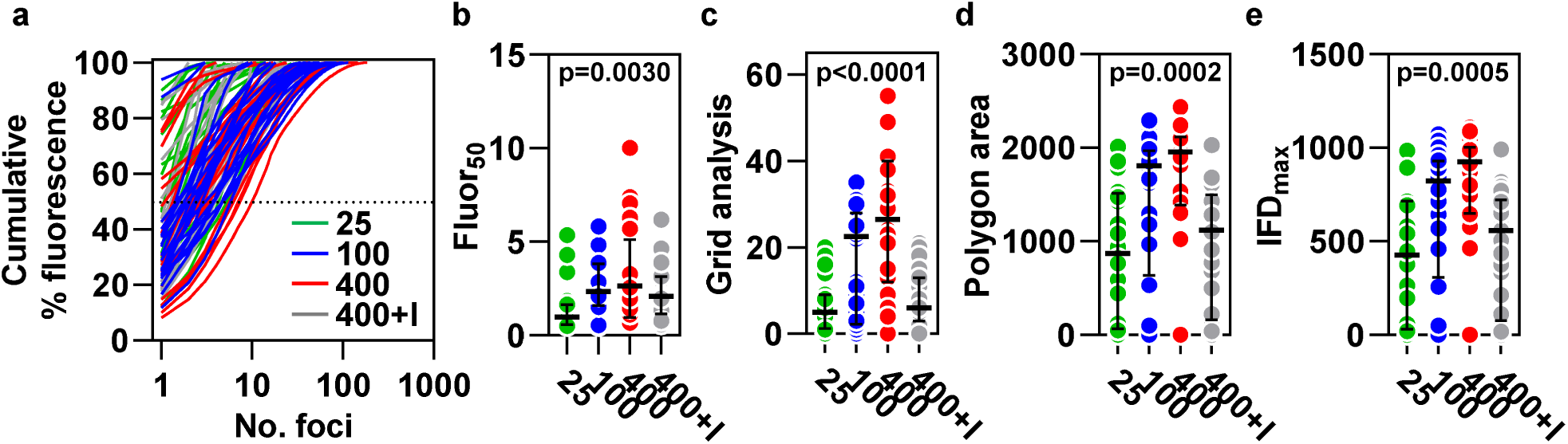
Quantitation of bacterial dissemination in response to a dose titration of *Mycobacterium marinum* (Mm) infection. Measures of dissemination in zebrafish larvae 4 dpi with 25, 100 or 400 cfu Mm ± 400 µM isoniazid (I) (n=24, 28, 22 and 25, respectively). **a**, The relationship between cumulative percentage fluorescence and the number of bacterial foci responsible for this signal. **b**, Fluor_50_, calculated by interpolation of the data shown in **a**. The three spatial dissemination parameters, **c**, grid analysis, **d**, polygon area and **e**, IFD_max_. Lines, **a**, and data points, **b-e**, represent individual zebrafish larvae. Lines and error bars, **b-e**, represent median ± IQR. p values were derived from Kruskal-Wallis tests with Benjamini, Krieger and Yekutieli correction for multiple comparisons.

We next quantified the relationship between our dissemination parameters and existing outcome measurements, by Spearman rank correlation analysis (Supplementary figure 5). None of our novel measures correlated perfectly with integrated fluorescence or the number of foci, which were highly correlated with each other. Fluor_50_, in particular, was less strongly associated with either of the existing measures of outcome compared to the other dissemination measurements. Grid analysis was highly correlated with the number of foci and polygon area almost perfectly correlated with IFD_max_. This is in keeping with the strategies used to derive these parameters, where the second variable is related to the first. Overall, this analysis suggests that our parameters represent distinct measures of outcome that likely reflect different aspects of infection.

We have generated a suite of tools to quantify spatial distribution of pathology and the proportional distribution of the total disease burden between sites in zebrafish larvae. In the context of infection, these novel measurements of outcome may provide greater resolution to detect differences following experimental manipulation, that might not otherwise be elucidated when principally reliant on pathogen burden as an outcome measure. This, in turn, could potentially provide new insights into immunological mechanisms of protection and pathogenesis or identify pathways for therapeutic intervention. We anticipate that our tools could also be applied to other zebrafish models where spatial distribution of focal pathology is important, such as metastatic cancer. As such, these analysis methods represent an exciting advance, of considerable utility to a wide range of investigators engaged in zebrafish research.

## METHODS

### ZEBRAFISH

Zebrafish were raised and maintained on a 14/10 light/dark cycle at 28.5°C according to standard protocols^5^ in the Zebrafish Facility at University College London. Work was approved by the British Home Office (Project License 70/8900). All experiments were conducted on larvae up to five days post fertilisation, before they are protected under the Animals (Scientific Procedures) Act. Adult AB/TL (wild type) zebrafish were spawned to generate embryos for infection experiments. Embryos were maintained at 28.5°C in egg water containing 60 µg / ml Tropic Marin Sea Salt (Norwood Aquarium) and anaesthetised with egg water containing 200 µg / ml buffered tricaine (3-aminobenzoic acid ethyl ester) (Sigma-Aldrich) during bacterial injections and imaging. Egg water was supplemented with 0.003% PTU (1-phenyl-2-thiourea) (Sigma-Aldrich) after bacterial injections had been performed, in order to inhibit melanisation.

### M. MARINUM INFECTION

*Mycobacterium marinum* (Mm) M strain expressing the pmsp12 mWasabi vector^4^ was cultured on Middlebrook 7H10 agar (Becton Dickinson and Company) supplemented with 0.5% oleic acid/albumin/dextrose/catalase (OADC) (Becton Dickinson and Company), 0.5% glycerol (Sigma-Aldrich) and hygromycin (50µg/ml) (Fisher Scientific). To generate injection inocula, Mm from agar plates was cultured statically at 28.5°C for 24 hours in Middlebrook 7H9 broth (Becton Dickinson and Company) supplemented with 10% albumin/dextrose/catalase (ADC) (Becton Dickinson and Company) and hygromycin (50µg/ml) (Fisher Scientific) and harvested at mid-log growth (optical density (OD)_600nm_ 0.7-1), (OD)_600nm_ 1 representing 10^8^ colony forming units (cfu) / ml. Harvested bacteria were washed three times in phosphate buffered saline (PBS) (Gibco) and resuspended in 2% polyvinylpyrrolidone (PVP)40 (Sigma-Aldrich)/PBS containing 10% phenol red (Sigma-Aldrich) to aid visualisation of injections^6^.

Zebrafish embryos staged at 24 hours post-fertilisation (hpf) were manually dechorionated using jeweller’s forceps (Dumont #5, World Precision Instruments), then infected at 28-30 hpf by caudal vein injection of Mm^6^. In three independent experiments from which images were used to develop our dissemination analysis tools, embryos were infected with 200-400 cfu Mm in a volume of 1 nl. For the Mm dose titration experiment, larvae were intravenously infected with 1 nl of bacterial suspension containing 25, 100 or 400 cfu. A subset of larvae infected with 400 cfu Mm were maintained in egg water containing 400 µM isoniazid (Sigma-Aldrich) to control infection^7^.

### STEREOFLUORESCENCE MICROSCOPY

Live, anaesthetised larvae were imaged four days post-infection (dpi) on a flat agarose plate, using an M205FA stereofluorescence microscope (Leica) with a 1x objective. Brightfield and fluorescence images were captured using a DFC365 FX camera (Leica) and exported as 8- or 16-bit TIF files for analysis.

### CLASSIFICATION OF IMAGES USED TO DEVELOP ANALYSIS TOOLS

To develop our analysis tools we used images of Mm infected zebrafish larvae in which fluorescence was represented as a binary channel to limit the visual impact of variable signal intensity. Images that were concordantly classified by subjective visual evaluation as representative of minimally, moderately or widely disseminated infection, by two independent investigators blinded to the experimental groups, were included.

### IMAGE ANALYSIS

Images were analysed using QuantiFish and a published ImageJ macro^4^ to quantify bacterial burden and number of bacterial foci. Analysis tools to quantify dissemination of bacterial infection were developed in Python 3, then integrated into QuantiFish, which was used to quantify dissemination of bacterial infection.

### QUANTIFISH

QuantiFish is an open source application written in Python 3 and released under the GNU General Public License (version 3). Compiled installers are available for Windows and Mac, and the source code can be run on other operating systems. QuantiFish utilises the SciPy (http://www.scipy.org), NumPy^8^, Pillow (https://github.com/python-pillow), and scikit-image^9^ libraries. The software provides an intuitive graphical user interface to enable automated analysis of fluorescence in zebrafish embryos. Images of individual fish in Tagged Image Format (.tif) are imported using the Python Imaging Library (Pillow fork) before being converted into NumPy arrays, which enables efficient manipulation by considering the image as an array of numbers. Values below the user-defined threshold are then removed to allow quantitation of the number of positive pixels and integrated fluorescence (the sum of all positive pixels) as surrogate measures of the total burden of pathology.

#### Dissemination analysis tools

To generate the dissemination measurements, detailed information from each individual area of fluorescent signal, termed a “focus”, must first be collected. Foci are detected by using scikit-image functions to identify pixels representing the peak intensity of an area of fluorescence and label continuous regions of signal above the desired threshold. The NumPy “unique” function is then used to evaluate the size of each labelled region before objects smaller than the user-defined minimum are excluded from further analysis. Objects are not further segmented due to the limited ability to separate touching features using a 2D image of a 3D embryo, alongside the limited resolution of images of entire fish, which prohibits visualisation of individual bacteria. Detected regions therefore represent foci of infection rather than individual cells or bacteria. The scikit-image “region properties” function is used to calculate and store an array of statistics for each region of fluorescence. The area, centre coordinates, minimum, maximum and mean intensity for each individual focus are extracted from the “region properties” array and optionally logged into a separate output file. Integrated intensity for each focus is calculated by multiplying the area of the focus by the mean intensity.

#### Fluor_50_

To derive the fluor_50_ statistic, the integrated intensity of foci in each image is ranked largest to smallest, the percentage of the total fluorescence within each focus is calculated, then the cumulative percentage intensity is determined using the NumPy “cumulative sum” function. This is plotted against the number of foci responsible for the signal, then the number of objects that contribute 50% of the total fluorescence (fluor_50_) is estimated by linear interpolation using SciPy interpolation classes.

#### Spatial distribution analysis parameters

For the spatial analyses, individual foci are considered as single points (centroids) to minimise skewing of data from unusually large objects. A list of focus centroid coordinates is extracted from the previously obtained “region properties” for each object and entered into an empty Boolean array in the shape of the original image, creating a “map” of centroids.

##### Grid analysis

To perform the “grid analysis” NumPy’s “array splitting” function divides the centroid map into squares of a size specified by the user. The algorithm classifies positive grid zones as those that contain coordinates for the centroid of any focus (designated “True” in the Boolean array), then quantifies the number of positive zones (as well as the total number of grid sections).

##### Polygon area

The list of focus centroid coordinates is also used to generate the “polygon area” parameter. We used the SciPy “ConvexHull” class to calculate the area of a polygon which encompasses the centroids of all foci, known as a convex hull.

##### Maximum inter-focus distance (IFD_max_)

Using the SciPy “Euclidean distance” function, the subset of points used to construct the polygon edges and vertices are taken forward to generate a distance matrix between all possible pairs of these centroids only, from which the two most widely separated are used to calculate the maximum inter-focus distance (IFD_max_).

##### Remark 1

Given that the centroids used to construct the polygon boundaries will always contain the two most distant points, restricting this analysis to these pairs of coordinates minimises the computational power needed to determine the IFD_max_. This is particularly relevant in images with large numbers of foci, for which analysis of all possible coordinate pairs would require substantially increased computational time.

##### Remark 2

It should be noted that a polygon cannot be generated in images containing fewer than three foci or where coordinates align to produce a 1D line with an area of zero. In these scenarios the software defaults to evaluation of all possible coordinate pairs to determine IFD_max_. However, it is extremely unlikely that images containing large numbers of foci would fail to produce a 2D polygon, hence the low probability of evaluating excessive numbers of irrelevant coordinate pairs that would lead to significantly increased computational time.

##### Output files

Results are written to two .csv files, one containing summary statistics for each image and a second, optional file containing data for each individual region of signal. Key settings such as the threshold used and the fluorescence channel analysed (in multi-channel images) are also logged.

### IMAGEJ MACRO

Images were also analysed using a published ImageJ macro as previously described^4^. This generates output data for the number of foci (“Count”), area of signal (“Total Area”), the percentage of the total area occupied by detected signal (“%Area”) and the average size of foci (“Average Size”). The number of positive pixels was calculated by multiplying “%Area” by the total number of pixels for each image.

### COMPARISON OF QUANTIFISH AND IMAGEJ

QuantiFish offers several significant improvements on existing methods to analyse fluorescence in zebrafish embryos. Most notably, in addition to the currently used statistics of integrated fluorescence (or pixel counts) and object counts, this software provides new measurements for the spatial dispersion of foci and for the distribution of pathology across sites, as a proportion of the total disease burden. These parameters were successfully applied to quantify dissemination of infection, providing novel measurements of outcome that we anticipate will have significantly greater resolution to detect differences compared to existing tools.

Performance was prioritised by using simple and efficient algorithms, ensuring that the analysis can be implemented rapidly, with minimal user input and without the need for dedicated hardware. The program utilises an intuitive user interface, with features such as automatic image bit depth detection which allows quantitation to be performed without the need for significant prior experience in image analysis. The program also supports selective filtering of files for analysis, eliminating the need for images to be manually sorted prior to quantitation. The inclusion of a previewing feature allows the user to visually inspect detected fluorescence in real time while configuring detection parameters, which facilitates the process of determining an appropriate threshold for a given data set. Overall, this software introduces novel dimensions for the analysis of fluorescence dissemination in zebrafish larvae while providing a simple and convenient interface which is accessible across the field.

## CODE AVAILABILITY

The source code and manual are available at https://github.com/DavidStirling/QuantiFish. The version of QuantiFish used in this paper is available as Supplementary Software.

## DATA AVAILABILITY

The data presented in this study are available from the corresponding author on request.

This work was supported by a UK Medical Research Council (MRC) fellowship to G.T. (MR/N007727/1), a Wellcome Trust Investigator Award to M.N. (207511/Z/17/Z), a Sir Henry Dale Fellowship jointly funded by the Wellcome Trust and the Royal Society to P.M.E. (105570/Z/14/Z) and the NIHR Biomedical Research Centre Funding to University College Hospitals NHS Foundation Trust and University College London.

## AUTHOR CONTRIBUTIONS

G.T. and M.N. conceived the idea. G.T., D.S., and O.S. performed the experiments. G.T. and P.M.E. classified the images. D.S. wrote the software. G.T., D.S., E.G. and M.N. conceived the analysis strategies. G.T., D.S. and V.T. analysed the data. G.T., D.S. and M.N. wrote the manuscript with input from all authors.

