## Supplementary figures for "Analysis tools to quantify dissemination of pathology in zebrafish larvae"

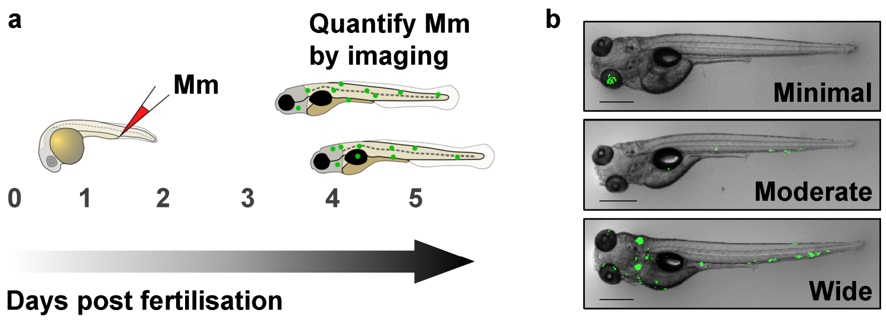


**Supplementary Figure 1: Infection of zebrafish larvae with *Mycobacterium marinum* (Mm).**

(a) Schematic diagram of the experimental design. Zebrafish embryos were infected at 28-30 hours post fertilisation (hpf) by caudal vein injection of *Mycobacterium marinum* (Mm) expressing mWasabi (green foci), then imaged by stereofluorescence microscopy after 4 days, to quantify bacterial burden, number of bacterial foci and dissemination. (b) Representative images of zebrafish larvae with minimally, moderately and widely disseminated Mm infection. Scale bars, 500 µm.

a b

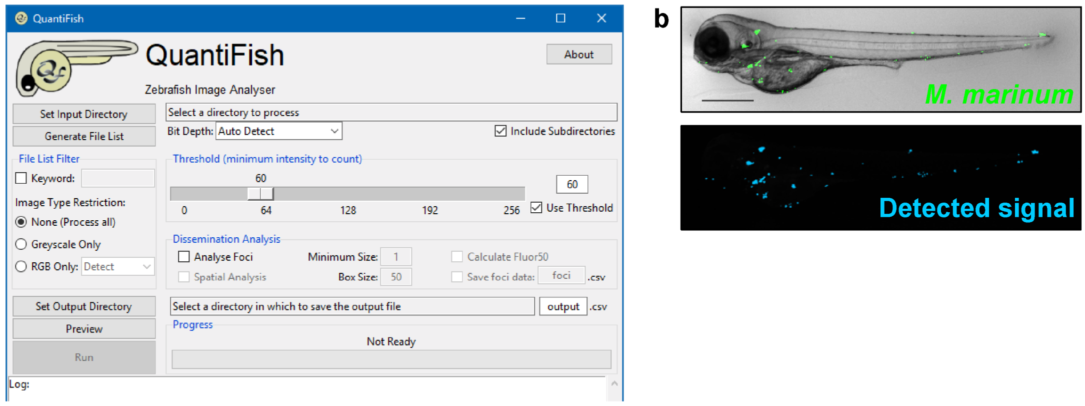


**Supplementary Figure 2: Detection and quantitation of fluorescence using QuantiFish.**

(a) QuantiFish offers an accessible interface with automatic image bit depth detection, selective file filtering and a previewing pane to visualise detected fluorescence, for rapid quantitation of bacterial burden, number of bacterial foci and four parameters of dissemination. (b) Bacterial fluorescence in a zebrafish larva 4 days post infection (dpi) with *Mycobacterium marinum* expressing mWasabi (upper panel), detected using QuantiFish (lower panel). Scale bar, 500 µm.

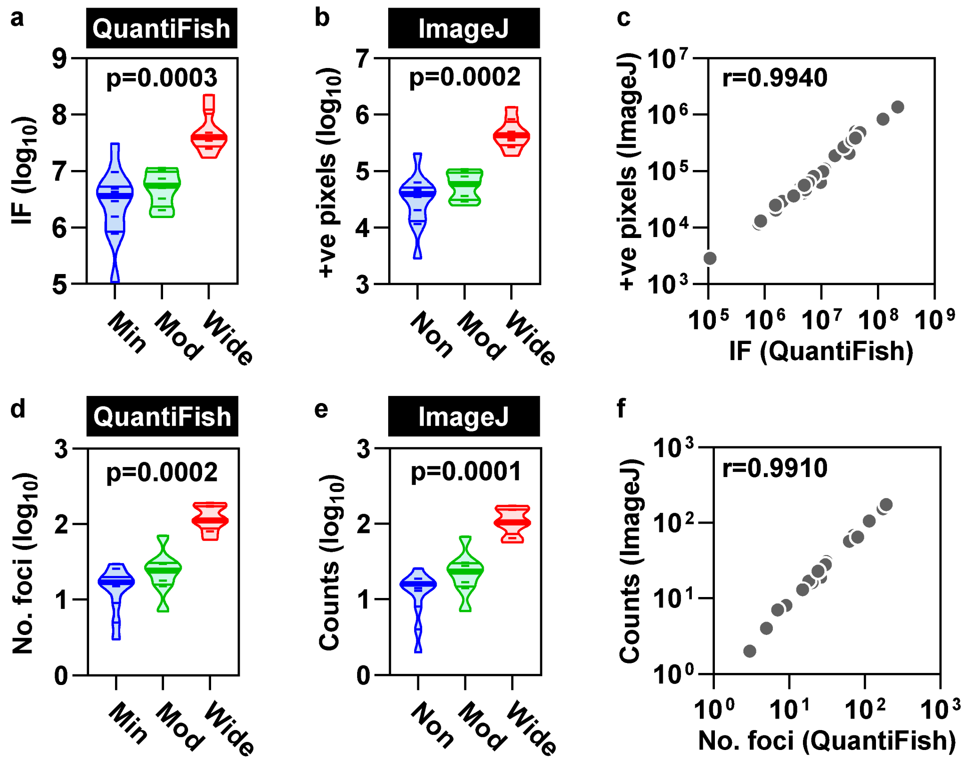


**Supplementary Figure 3: Equivalent performance of QuantiFish and ImageJ for quantitation of bacterial burden and counting bacterial foci.**

(a) Integrated fluorescence (IF) detected using QuantiFish and (b) pixel counts determined using ImageJ, as surrogate measures of total bacterial burden in zebrafish larvae with minimally, moderately and widely disseminated *Mycobacterium marinum* infection (n=11, 8 and 8, respectively). (c) The relationship between IF and pixel counts. (d) The number of bacterial foci detected using QuantiFish and (e) ImageJ and (f) the relationship between these measurements. In violin plots (a, b, d, e) short dashes represent data points for individual zebrafish larvae and horizontal bars represent median ± IQR. Data points (c, f) represent individual zebrafish larvae. p values were derived from Kruskal-Wallis tests with Benjamini, Krieger and Yekutieli correction for multiple comparisons. r values were derived from Spearman rank correlation tests.

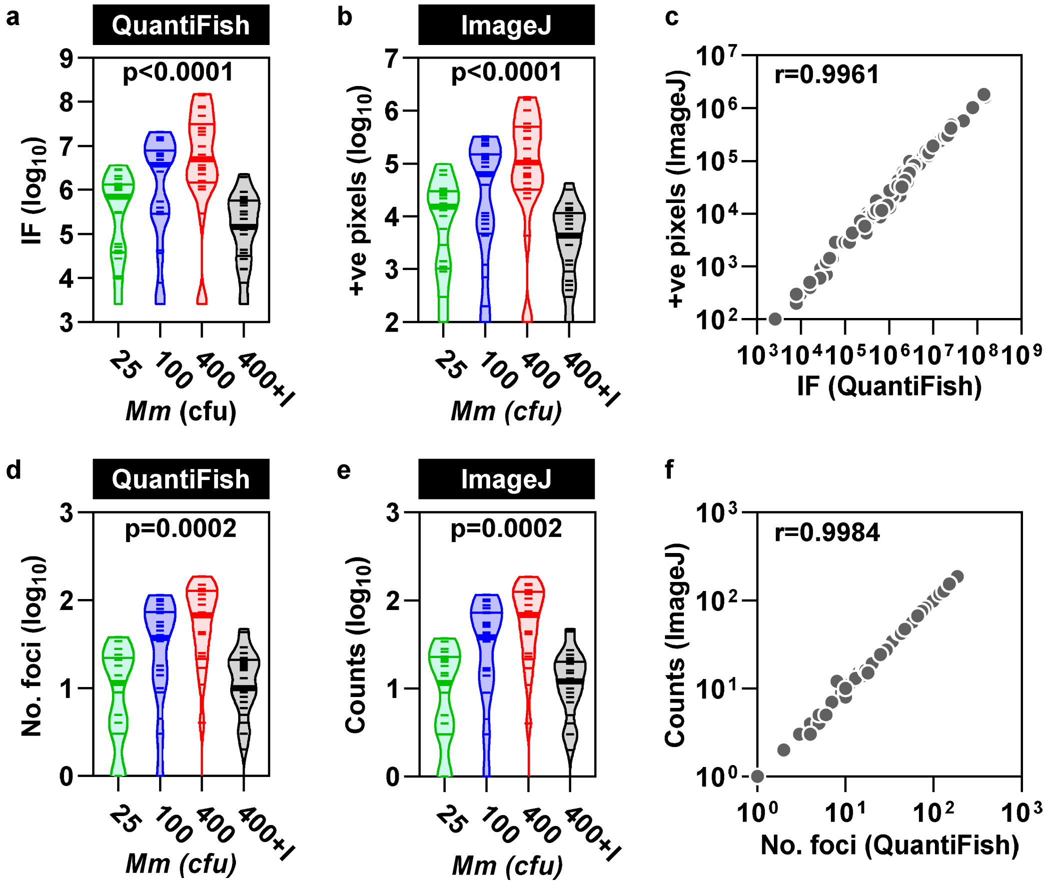


**Supplementary Figure 4: Quantitation of bacterial burden and number of bacterial foci in response to a dose titration of *Mycobacterium marinum* (Mm) infection.**

(a) Integrated fluorescence (IF) detected using QuantiFish and (b) pixel counts determined using ImageJ, as surrogate measures of total bacterial burden in zebrafish larvae 4 dpi with 25, 100 or 400 cfu Mm ± 400 μM isoniazid (I) (n=24, 28, 22 and 25, respectively). (c) The relationship between IF and pixel counts. (d) The number of bacterial foci detected using QuantiFish and (e) ImageJ and (f) the relationship between these measurements. In violin plots (a, b, d, e) short dashes represent data points for individual zebrafish larvae and horizontal bars represent median ± IQR. Data points (c, f) represent individual zebrafish larvae. p values were derived from Kruskal-Wallis tests with Benjamini, Krieger and Yekutieli correction for multiple comparisons. r values were derived from Spearman rank correlation tests.

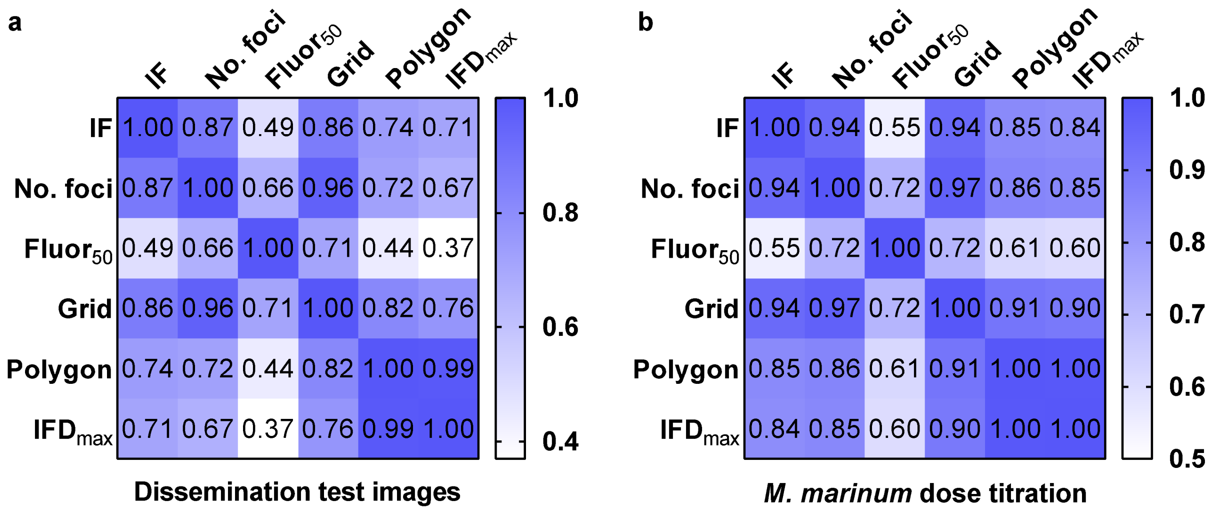


**Supplementary Figure 5: Correlation matrix of relationships between integrated fluorescence, number of foci and parameters of dissemination.**

Spearman rank correlation matrices of the associations between existing measurements of outcome of bacterial infection; integrated fluorescence (IF) and number of foci and our new measures of dissemination; $fluor_{50}$, grid analysis (Grid), polygon area (Polygon) and $IFD_{max}$ in (a) the images of *M. marinum* (Mm) infected larvae used to develop our analysis tools (n=27) and (b) the Mm dose titration experiment (n=99).
